# Progranulin deficiency leads to reduced glucocerebrosidase activity

**DOI:** 10.1101/540450

**Authors:** Xiaolai Zhou, Daniel H. Paushter, Mitchell D. Pagan, Dongsung Kim, Raquel L. Lieberman, Herman S. Overkleeft, Ying Sun, Marcus B. Smolka, Fenghua Hu

## Abstract

Mutation in the *GRN* gene, encoding the progranulin (PGRN) protein, shows a dose-dependent disease correlation, wherein haploinsufficiency results in frontotemporal lobar degeneration (FTLD) and complete loss results in neuronal ceroid lipofuscinosis (NCL). Although the exact function of PGRN is unknown, it has been increasingly implicated in lysosomal physiology. Here we report that PGRN interacts with the lysosomal enzyme, glucocerebrosidase (GBA), and is essential for proper GBA activity. GBA activity is significantly reduced in tissue lysates from PGRN-deficient mice. This is further evidence that reduced lysosomal hydrolase activity may be a pathological mechanism in cases of GRN-related FTLD and NCL.

## Introduction

Progranulin (PGRN), encoded by the *GRN* gene in humans, is a glycoprotein comprised of 7.5 conserved and highly disulfide-bonded homologous granulin domains connected by short linker regions (1-6). While the exact function of PGRN remains elusive, it has been found to be involved in numerous normal physiologic and pathologic processes, including regulation of inflammation, wound healing, and tumorigenesis, and it has also been shown to function as a growth and neurotrophic factor (7-16). Mutation in *GRN* has been linked to two neurodegenerative diseases with dose-dependent associations. Heterozygous mutation, resulting in PGRN haploinsufficiency, is known to cause frontotemporal lobar degeneration (FTLD), a clinically and pathologically heterogeneous disease often resulting in early-onset dementia (17-23). Homozygous mutation, resulting in complete loss of PGRN, causes neuronal ceroid lipofuscinosis (NCL), a lysosomal storage disease (24, 25). Although these two diseases are distinct, NCL-related phenotypes have been reported in FTLD patients with *GRN* mutation, suggesting that lysosomal dysfunction might serve as a common mechanism (26-28).

In addition to the connection to NCL, increasing lines of evidence indicate that PGRN is likely to be involved in lysosomal physiology. In addition to being lysosomally localized, having two independent trafficking pathways to the lysosome (29, 30), and being under transcriptional regulation with the majority of known lysosomal proteins (31), PGRN has recently been shown to be proteolytically processed to produce stable and functional granulin peptides in the lysosome (32-34), and is linked to the direct or indirect regulation of two lysosomal hydrolases, cathepsin D (CTSD) and glucocerebrosidase (GBA). PGRN was shown to stabilize and directly modulate the activity of cathepsin D *in vitro* and PGRN deficiency results in a reduction of cathepsin D activity *in vivo* (28, 35, 36). In another study, PGRN was purported to be a co-chaperone of glucocerebrosidase (GBA) (37, 38), a β-glucosidase mutated in Gaucher disease, in tandem with heat shock protein 70 (HSP70) (39). *Grn* knockout mice with induced chronic inflammation display cellular phenotypes similar to those with Gaucher disease (37, 38).

To better understand the lysosomal role of PGRN, we performed a stable isotope labeling by amino acids in cell culture (SILAC)-based proteomic screen for PGRN protein interactors. Corresponding with the previous findings, one of the top hits was GBA. In this study, we demonstrate that PGRN loss results in a substantial decrease in GBA activity in mouse tissues without any changes in protein levels.

## Material and Methods

### Primary Antibodies and Reagents

The following antibodies were used in this study: M2 mouse anti-FLAG (Sigma), M2 mouse anti-FLAG-conjugated beads (Sigma), mouse anti-myc (9E10), mouse anti-GAPDH (Proteintech Group), mouse anti-GBA (MilliporeSigma), rat anti-mouse LAMP1 (BD Biosciences), and sheep anti-mouse PGRN (R&D Systems). Rabbit anti-mouse PSAP and PGRN antibodies were produced as previously described (30), GFP-Trap beads (ChromoTek). 4-Methylumbelliferyl β-D-glucopyranoside (4-MU) GBA substrate (Sigma).

### Plasmids

Human GBA cDNA in the pDONR223 vector from the human ORFeome 8.1 collection was a gift from Dr. Haiyuan Yu (Cornell University). GBA was cloned into pcDNA3.1/myc-His A vector (Thermo Fisher Scientific) after digestion with EcoRI and XhoI. Human PGRN in the pCMV-Sport6 vector was obtained as previously described (29). GFP-PGRN was produced as previously described (30). GFP-granulin peptides were produced as described (36).

### Protein Production and Purification

GST-granulin E, GST-granulin F and His-SUMO-saposin C proteins were expressed from Origami B(DE3) bacterial strains (MilliporeSigma) with 0.1 mM IPTG induction overnight at 18°C. Proteins were purified using GST or cobalt beads. His-PGRN was purified with cobalt beads from the culture media of HEK293T cells as previously described (30). All purified proteins were concentrated and transitioned to PBS buffer with Centricon Centrifugal Filter Units (MilliporeSigma).

### Mouse Strains

C57/BL6 and *Grn^-/-^* mice (40) were obtained from The Jackson Laboratory. *Gba^-/-^* mice were generated as previously described (41). Mixed male and female mice were used for this study.

### Cell Culture

HEK293T and BV2 cells were maintained in Dulbecco’s Modified Eagle’s Medium (Cellgro) supplemented with 10% fetal bovine serum (Gibco) in a humidified incubator at 37°C with 5% CO2. WT, *Grn^-/-^* and *Gba^-/-^* mouse fibroblasts were cultured as described (30).

### Transfection, Immunoprecipitation, and Western Blot Analysis

Cells were transfected with polyethylenimine as previously described (42). Cells were lysed in a cold, near-neutral pH solution containing 150 mM NaCl, 50 mM Tris pH 7.5 or 50mM sodium acetate pH5.3, 1% Triton X-100, 0.1% deoxycholic acid, 1X protease inhibitors (Roche). After centrifugation at 14,000 xg, for 15 minutes, at 4°C, supernatants were transferred to clean tubes on ice, to which rabbit anti-PGRN antibody-conjugated Affi-Gel 15 (Bio-Rad Laboratories) or GFP-Trap beads were added, then rocked for 3-4 hours at 4°C. Samples were run on 12% polyacrylamide gels or 4-12% Bis-Tris gels (Invitrogen), then transferred to Immobilon-FL polyvinylidene fluoride membranes (Millipore Corporation) or nitrocellulose membranes (Millipore Corporation). Membranes were blocked with either 5% non-fat milk in PBS or Odyssey Blocking Buffer (LI-COR Biosciences) for 1 hour then washed with Tris-buffered saline with 0.1% Tween-20 (TBST) 3x for 5 minutes each. Membranes were incubated with primary antibodies, rocking overnight at 4°C, then washed as above, incubated with secondary antibodies for 2 hours at room temperature, then washed again. Membranes were scanned using an Odyssey Infrared Imaging System (LI-COR Biosciences). Densitometry was performed with Image Studio (LI-COR Biosciences).

### Tissue Preparation for Enzyme Assays and Western Blot

Tissues from 2-month-old WT and Grn^-/-^ mice were homogenized on ice with a glass Dounce homogenizer in a cold solution of either 1% (w/v) sodium taurocholate and 1% (v/v) Triton X-100, pH 5.2, for 4-MU activity assays or 0.2% (w/v) sodium taurocholate and 0.1% (v/v) Triton X-100 for MDW941 assays. Protein concentrations were determined via Bradford assay, then standardized.

### GBA Activity Assay with 4-MU Substrate

Tissue lysates, prepared as described, were diluted in a cold buffer of 0.1 M citric acid/0.2 M disodium phosphate (pH 5.0), with 2 mg/mL bovine serum albumin added. Ten microliters of each sample were added to 75 μL of cold 10 mM 4-MU substrate in the same buffer and incubated at 37°C for 30 minutes. Reactions were stopped by the addition of 200 μL of a 0.3 M glycine/0.2 M sodium carbonate (pH 10.7) stop solution. Plates were read at 360 nm excitation/460 nm emission with an Infinite M1000 microplate reader (Tecan). To test for the direct activation of recombinant GBA, a 50 μL reaction mixture containing 1 mg/mL BSA, 0.1 M sodium acetate (pH 4.5), 0.02 U Cerezyme, 0.2 mM 4-MU substrate, 0.1% Triton X-100, and 1, 5, or 9 μM recombinant PGRN, Grn E, Grn F, saposin C, GST, or an equal volume of PBS was incubated at 37°C for 30 minutes. Reactions were stopped by the addition of 50μL of a 0.32 M glycine/0.32 M sodium carbonate (pH 10.4) stop solution. Plates were read at 340 nm excitation/420 nm emission with an Infinite M1000 microplate reader (Tecan).

### Active GBA Assessment with MDW941 Probe

MDW941 was diluted to 100nM in tissue lysates, which were then incubated at 37°C for 30 minutes. Reactions were stopped by the addition of an equal volume of 2x Laemmli sample buffer with 10% β-Mercaptoethanol before heating at 95°C for 5 minutes. An equal amount of each sample (50μg total protein) was run on a 12% polyacrylamide gel, which was scanned at 532 nm excitation/580 nm emission with a Typhoon Imaging System (GE Healthcare), then western blot and assessment were performed as described above, with all values normalized to GAPDH.

### MDW941 Cell Labeling, Immunostaining, and Confocal Microscopy

WT and Grn^-/-^ MEFs were cultured on glass coverslips overnight. The next day, MDW941 was diluted to 5 nM in culture media, then equal volumes were added to each well and the plate was incubated at 37°C for 2 hours. Cells were washed 2x with PBS, fixed with 3.7% paraformaldehyde for 15 minutes at room temperature, followed by 3 additional PBS washes. Cells were permeabilized with Odyssey Blocking Buffer LI-COR Biosciences) + 0.05% saponin for 30 minutes at room temperature. Primary antibodies were diluted in the same buffer and added to coverslips, which were incubated in a humidified chamber overnight at 4°C. Coverslips were washed 3x with PBS, for 5 minutes each, then secondary antibodies diluted in the same blocking/permeabilization solution were added to the coverslips, which were incubated at room temperature, in the dark, for 2 hours. After 3 additional PBS washes, coverslips were mounted on slides with Fluoromount-G (SouthernBiotech). Images were acquired with a CSU-X series spinning disc confocal microscope (Intelligent Imaging Innovations) with an HQ2 CCD camera (Photometrics) using a 100x objective.

### SILAC and Mass Spectrometry

BV2 cells were infected with control lentivirus (pLenti-CRISPR2, Addgene) or lentivirus expressing Cas9 and guide RNA (sequence 5’-GCTCCCTGGGAGGCATCTGG-3’) and selected with puromycin. The cells were then grown a minimum of five generations in DMEM with 10% dialyzed fetal bovine serum (Sigma) supplemented with either heavy (C13, N15 arginine and lysine) amino acids or light (C12, N14 arginine and lysine). Cells were grown to confluency in two 15-cm plates, each, and lysed then immunoprecipitated as described above using homemade rabbit anti-PGRN antibodies bound to Affi-Gel 15 (Bio-Rad Laboratories). Samples were then mixed and boiled 5 minutes with 10mM DTT followed by alkylation by treating samples with a final concentration of 28 mM iodoacetamide. Proteins were precipitated on ice for 30 minutes with a mixture of 50% acetone/49.9% ethanol/0.1% acetic acid. Protein was pelleted and washed with this buffer, reprecipitated on ice, and dissolved in 8 M urea/50 mM Tris (pH 8.0) followed by dilution with three volumes of 50 mM Tris (pH 8.0)/150 mM NaCl. Proteins were digested overnight at 37°C with 1 μg of mass spectrometry grade Trypsin (Promega). The resulting peptide samples were cleaned up for mass spectrometry in a Sep-Pak C18 column (Waters) as follow: samples were acidified with 0.25% formic acid and 0.25% trifluoroacetic acid, loaded onto a preequilibrated C18 column, washed twice with 0.1% acetic acid and eluted with 80% acetonitrile/0.1% acetic acid into silanized vials (National Scientific). Samples were evaporated using a SpeedVac and then redissolved in 70% acetonitrile with ~1% formic acid. Peptides were separated using hydrophilic interaction liquid chromatography on an Ultimate 300 LC (Dionex). Each fraction was evaporated in a SpeedVac and resuspended in 0.1% trifluoroacetic acid with spiked-in angiotensin II as an internal standard. Samples were run on a QE Orbitrap XL mass spectrometer (Thermo Fisher Scientific). XPRESS software, part of the Trans-Proteomic Pipeline (Seattle Proteome Center), was used to quantify all the identified peptides. Mann-Whitney U test was used to calculate the p value to generate the volcano plot.

## Results

### PGRN interacts with GBA

SILAC was performed in the murine microglia-derived BV2 cell line, with either normal PGRN expression or loss of PGRN expression after CRISPR/cas9 mediated genome editing (43, 44) (Fig. 1a and 1b). GBA was one of the high-confidence hits for PGRN from this experiment (Fig. 1b) (Table S1).

**Figure 1.**
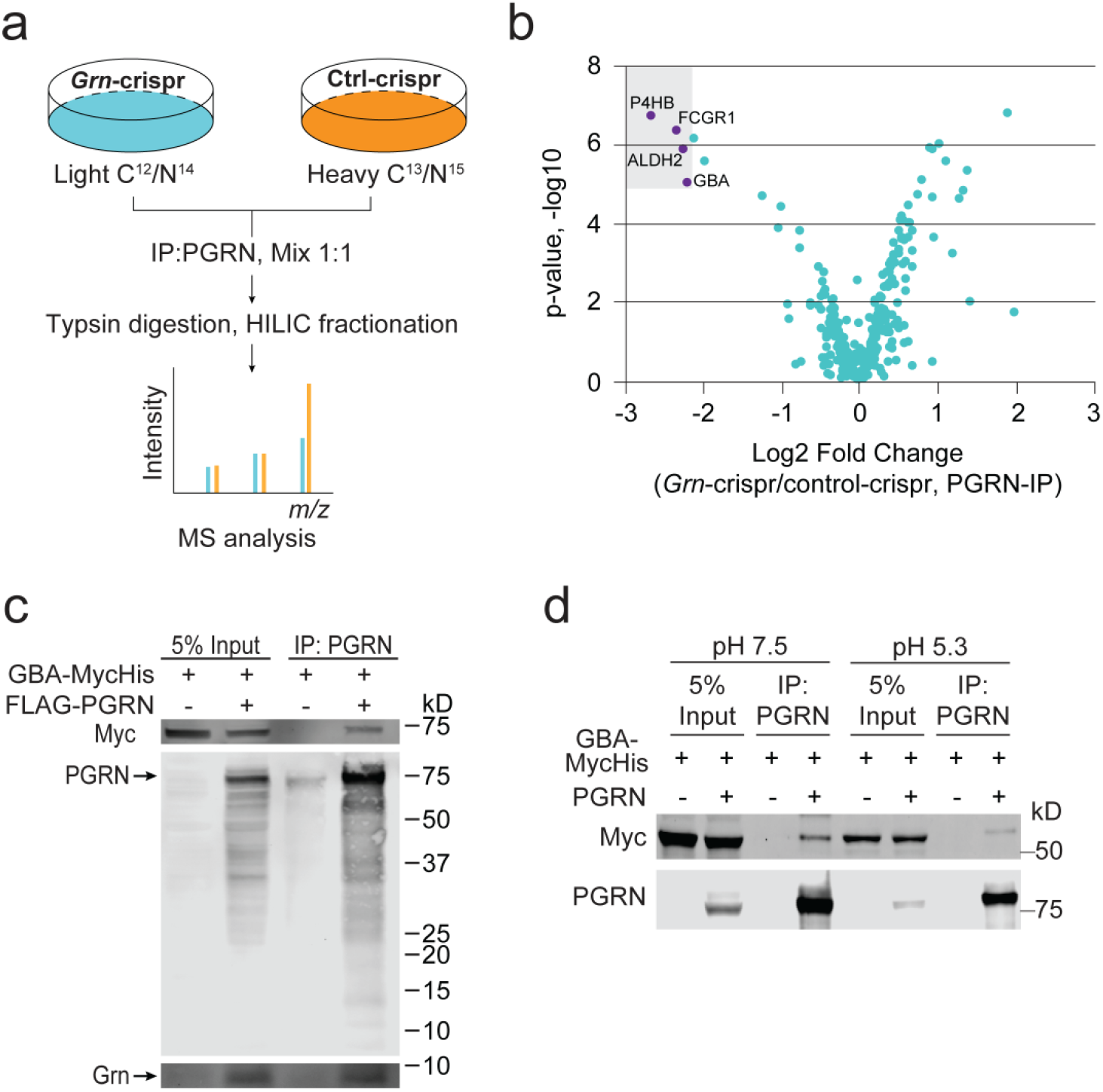
PGRN interacts with GBA. **(a)** Schematic illustration of the SILAC experiment searching for PGRN interactors. **(b)** Volcano plot of SILAC hits. Top hits identified in the heavy fraction are highlighted. **(c)** Anti-PGRN co-immunoprecipitation of FLAG-PGRN and GBA-myc overexpressed in HEK293T cells. **(d)** Anti-PGRN co-immunoprecipitation of PGRN and GBA-myc overexpressed in HEK293T cells using buffers of pH7.5 and pH5.3.

To verify the physical interaction between PGRN and GBA, FLAG-PGRN and GBA with a C-terminal myc tag were co-transfected in HEK293T cells, then anti-PGRN immunoprecipitation (IP) was performed. GBA-myc signal was detected in the IP products from the PGRN transfected, but not control samples, indicating a specific binding interaction (Fig. 1c). This interaction persists at pH5.3 (Fig. 1d). Additionally, PGRN has been shown to be processed into individual granulin peptides within the lysosome (32, 33) and two of these peptides, Grn E and Grn F, have been shown to interact with GBA (37, 38). Since both PGRN and Grn peptides were pulled down in the anti-PGRN IP (Fig. 1c), we tested whether GBA can bind specific Grn peptides. N-terminal GFP-tagged Grn peptides or GFP-PGRN were co-transfected with myc-tagged GBA, followed by anti-GFP IP. While there was no obvious binding of granulin E, granulin F did interact with GBA in the cell lysate, as well as granulin A, although to a lesser extent (Fig. 2a). However, it is not clear whether these granulin peptides are folded correctly when expressed individually. We repeated the experiments with secreted granulins, which, having proceeded through the entire secretory pathway, may be more likely to be properly folded. In this instance, all the granulins except Grn G shows weak binding to GBA (Fig. 2b).

**Figure 2:**
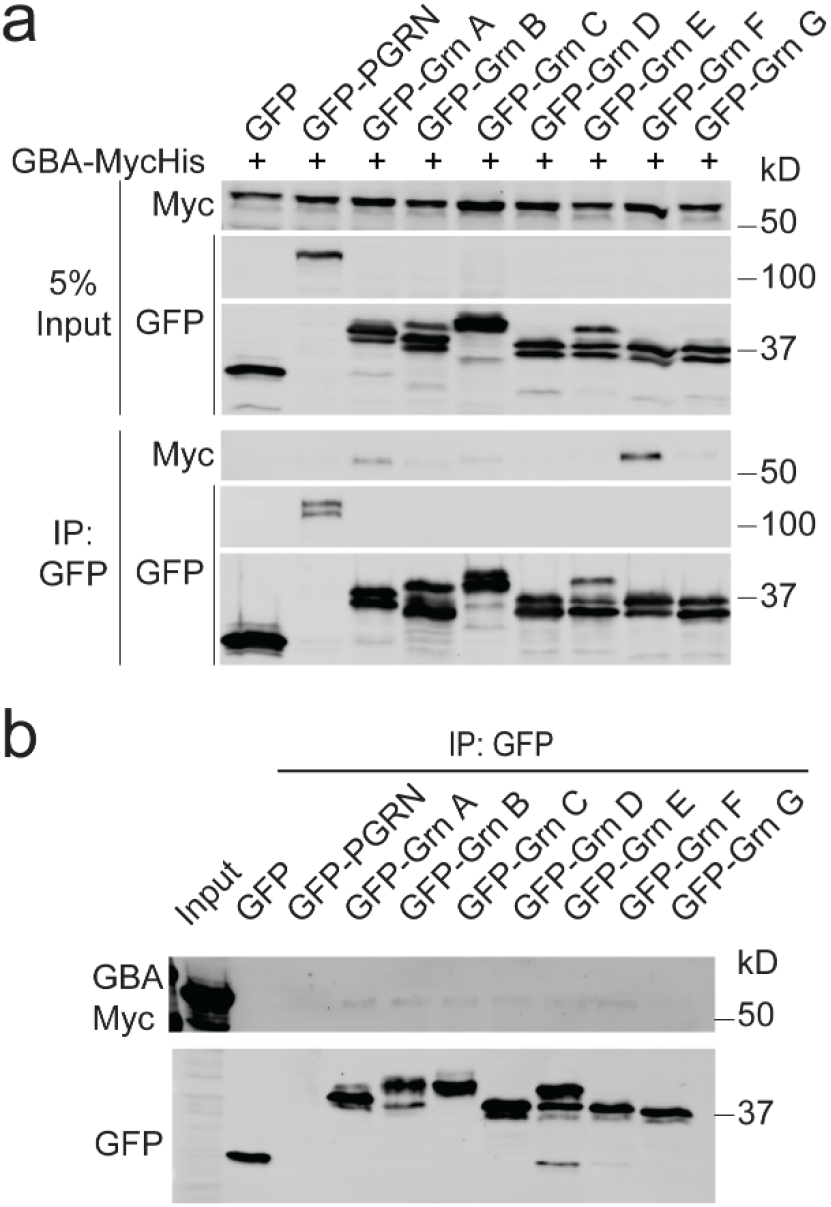
Interaction between GBA and granulin peptides. **(a)** Anti-GFP co-immunoprecipitation of GFP-PGRN or individual GFP-granulins and GBA-myc overexpressed in HEK293T cells shows binding primarily to Grn F, with weaker binding to Grn A and other Grn peptides. **(b)** Anti-GFP co-immunoprecipitation of GFP-PGRN or individual GFP-granulins and GBA-myc from conditioned medium of transfected cells shows weaker binding to all the granulins except Grn G.

### PGRN deficiency leads to reduced GBA activities

To determine whether PGRN regulates GBA activities, we performed an *in vitro* GBA activity assay using a well-established fluorogenic GBA substrate, 4-Methylumbelliferyl β-D-glucopyranoside (4MU-β-glc) (45-47) (Fig. 3). GBA activity was measured in tissue lysates from 2-month-old WT and *Grn*^-/-^ mice, before obvious lysosomal phenotypes were observed. Liver and spleen lysates from *Grn*^-/-^ mice showed a significant decrease in GBA activity compared to WT controls (Fig. 3a,3b). While the cerebrum and cerebellum showed a trend toward decreased GBA activity, it was not significant (Fig. 3c, 3d). However, the midbrain, where we tend to observe the most severe defects in our knockout mouse line, showed a significant decrease in GBA activity (Fig. 3e). Thus PGRN is likely to have cell type and tissue specific effects on GBA.

**Figure 3.**
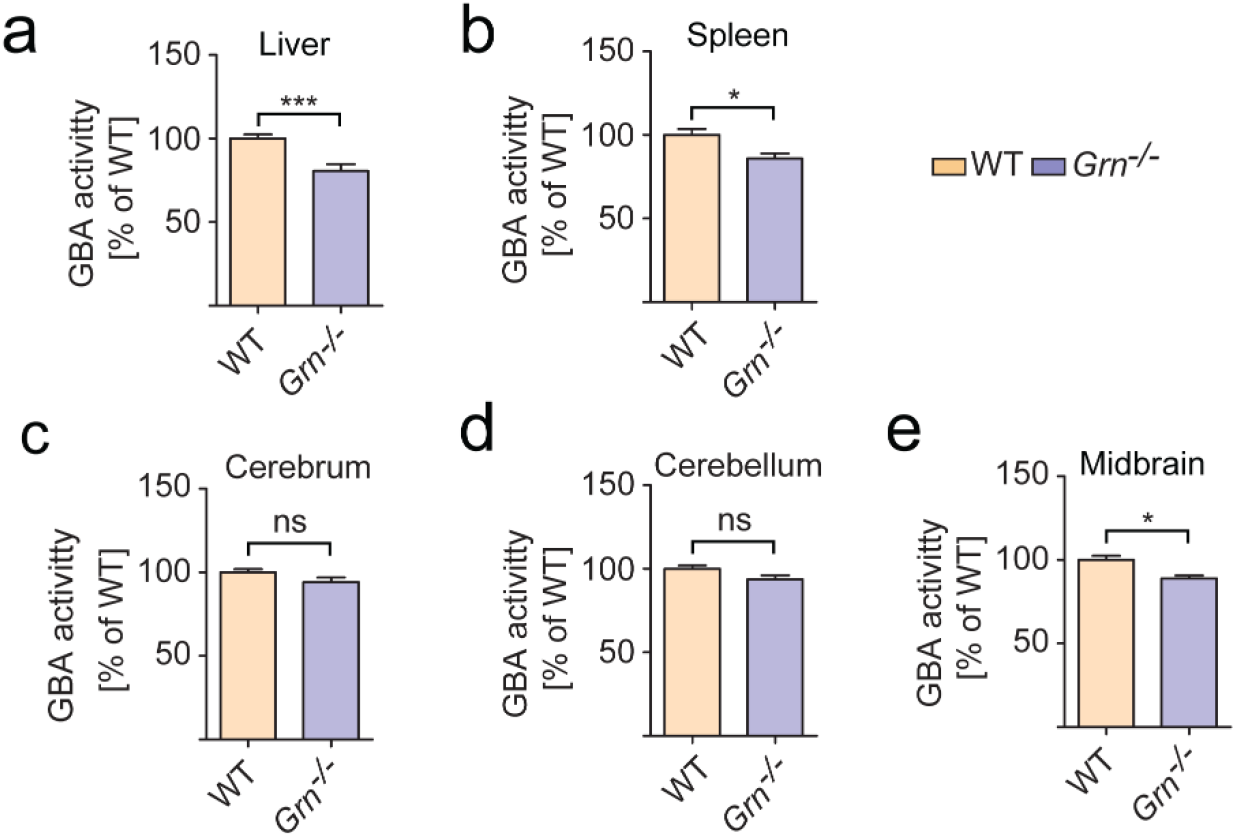
PGRN deficiency results in decreased GBA activity in mouse tissues. (GBA activity was assessed in tissue lysates from 2-month-old WT and *Grn*^-/-^ mice, as indicated, using the substrate 4-MU. (n = 5–6, ±SEM, *p value <0.05, **p-value <0.01, ns, not significant, Student’s t-test.

These results were confirmed using a recently developed ultrasensitive fluorescent probe, MDW941, which specifically reacts with active forms of the enzyme and has been shown to be sensitive enough to detect the activity of recombinant GBA in the attomolar range (48-50). Using this probe, an even greater disparity between GBA activity in WT and *Grn^-/-^* tissues was observed (Fig. 4a-4e). To verify that any changes in activity were not due to alterations in total GBA protein levels, SDS-PAGE and western blot of the tissue lysates were performed using antibodies specific to GBA (Fig. S1), with no significant differences seen between groups (Fig. 4a-4e).

**Figure 4.**
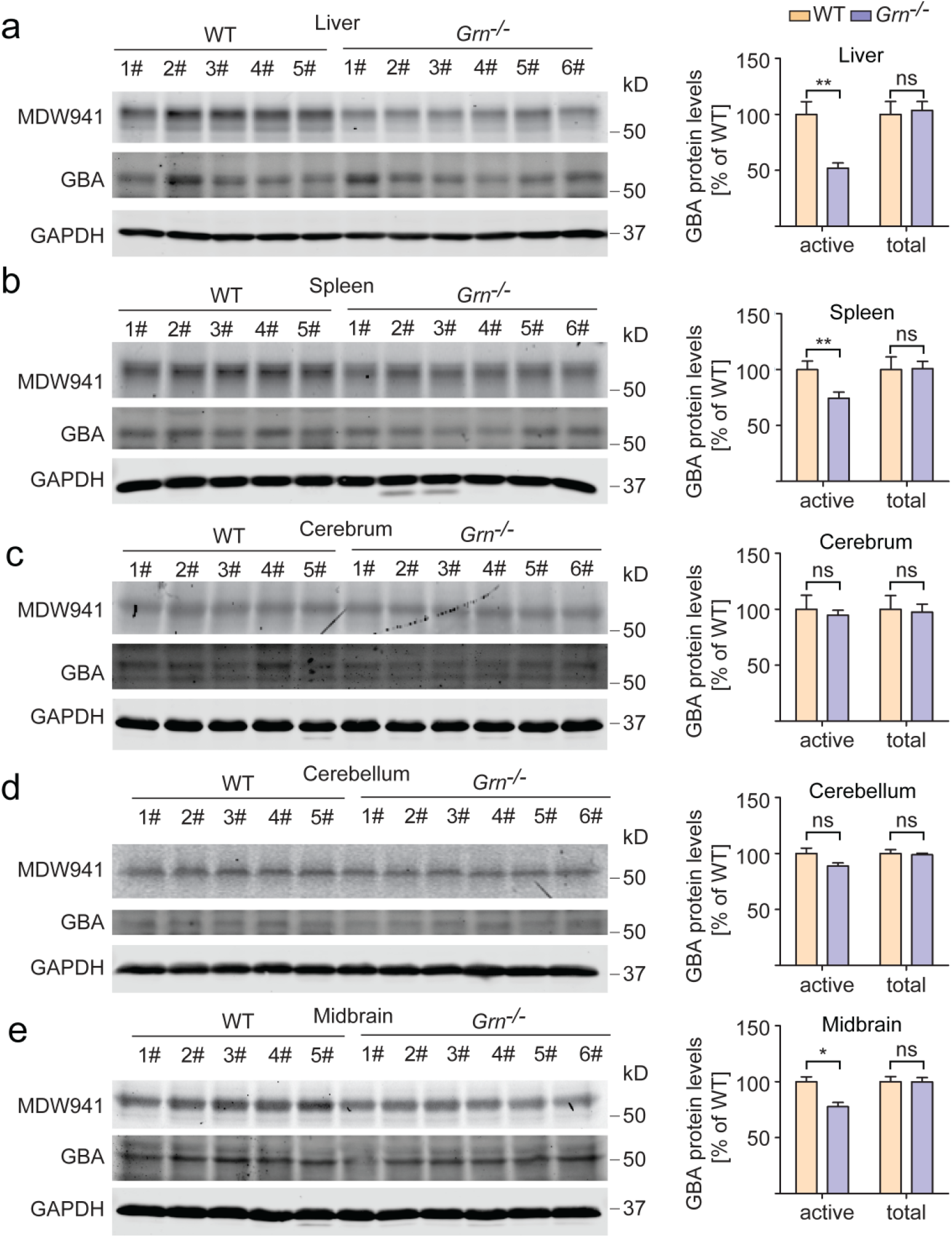
PGRN deficiency results in decreased GBA activity without changes in GBA protein levels. Tissue lysates from 2-month-old WT and *Grn^-/-^* mice were incubated with the MDW941 GBA activity probe. The samples were run on SDS-PAGE and MDW941 labeled GBA (active) was detected using a fluorescent scanner, then western blot was performed to assess total GBA protein levels. n = 5–6, ±SEM, *p value <0.05, **p-value <0.01, ns, not significant, Student’s t-test.

Previously we showed that PGRN binds to prosaposin (PSAP) to facilitate each other’s lysosomal trafficking (30, 51, 52). PSAP is known to get cleaved in the lysosome to generate saposin peptides, of which saposins A and C are known activators of GBA (46, 53, 54). This raises the possibility that the observed decrease in GBA activity with PGRN loss could be due to a decrease in PSAP or saposin peptides. However, we did not observe a significant change in PSAP or total saposins in the PGRN knockout tissues where GBA activity was reduced (Fig. 5). Similarly, no obvious difference in saposin C levels in WT and PGRN deficient liver lysates were detected following immunoprecipitations using antibodies specific for saposin C (Fig. S2). Unfortunately we cannot test saposin A levels due to a lack of specific antibodies.

**Figure 5.**
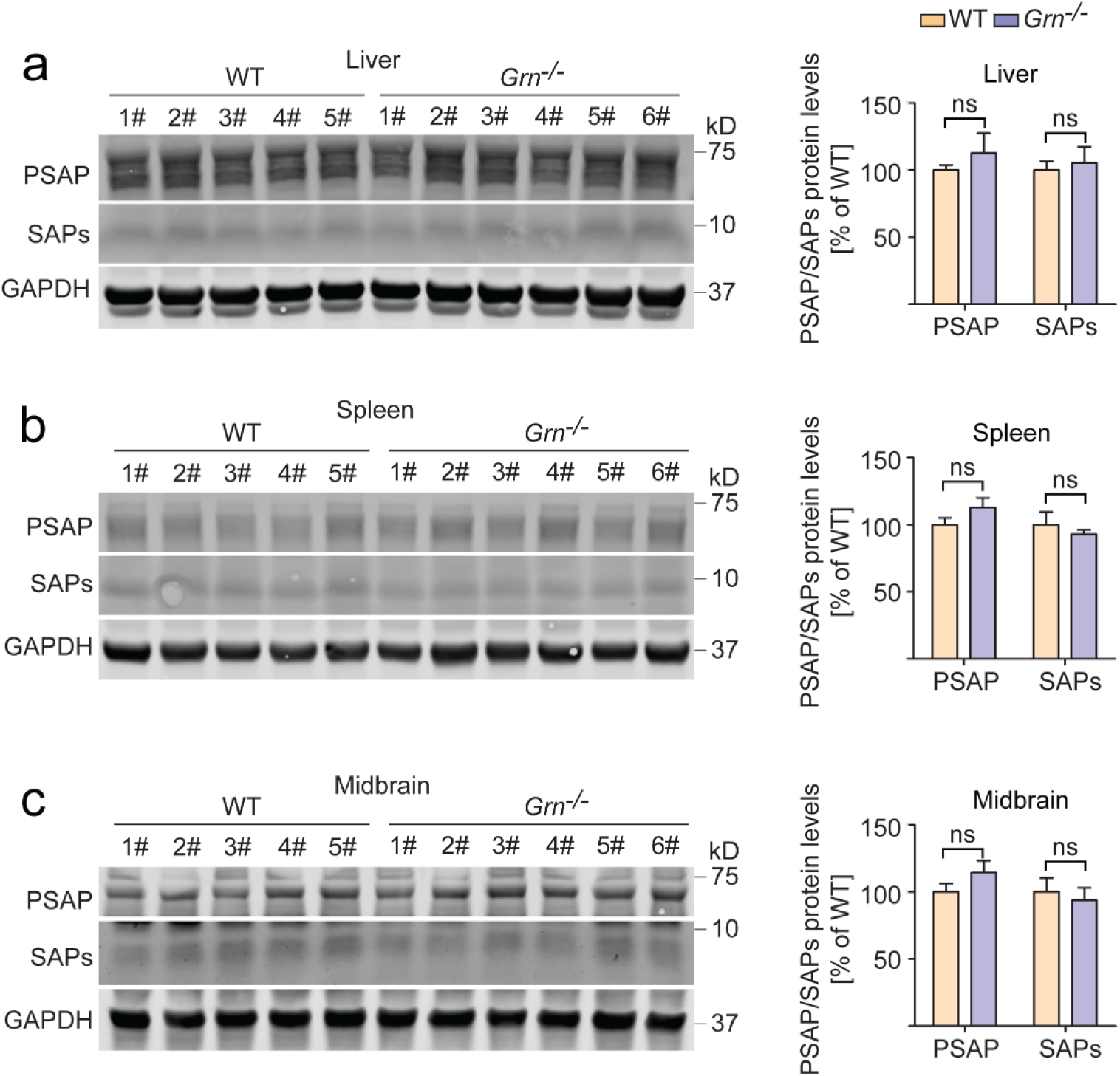
PSAP and total saposin levels are not changed in PGRN deficient mice. Mouse tissue lysates from 2-month-old WT and *Grn^-/-^* mice were assessed for PSAP and total saposin peptide levels, which were normalized to GAPDH. n = 5–6, ±SEM, ns, not significant, Student’s t-test.

With the changes we observed in GBA activity, and because PGRN and Grn E have been shown to directly activate CTSD *in vitro,* we wanted to test whether PGRN, Grn E, or Grn F could directly augment the activity of GBA. We performed a 4-MU activity assay of a pharmaceutical grade recombinant GBA (Cerezyme) with the addition of recombinant His-PGRN, GST-Grn E, GST-Grn F, His-SUMO-saposin C, or GST control. While the addition of saposin C greatly enhanced GBA activity, such effects were not observed with PGRN or granulin peptides (Fig. S3).

## Discussion

In this study, we demonstrate a physical interaction between the FTLD-related protein, PGRN, and the lysosomal enzyme, GBA, consistent with previous reports (37, 38). Additionally, among the granulin peptides, granulin F and to a lesser extent, granulin A, but not the previously reported granulin E, were shown to bind GBA in the cell lysate (Fig. 2a). All the secreted granulins except Grn G show weak binding to GBA (Fig. 2b).

The previous studies examining the relationship between PGRN and GBA primarily utilized a chronic inflammation model based on the administration of ovalbumin (OVA) to WT and *Grn^-/-^* mice over the course of multiple weeks. The authors showed that PGRN interacts with GBA and acts as a cochaperone of GBA and its trafficking receptor, lysosome membrane protein 2 (LIMP-2) (37, 38). Furthermore, LIMP-2 and GBA were shown to aggregate in the cytoplasm in PGRN deficient macrophages with experimentally induced chronic lung inflammation models (37, 38). Unfortunately, we were unable to find a single commercial antibody that specifically recognize the endogenous GBA protein in immunostaining using WT and *Gba^-/-^* mouse fibroblasts (data not shown), thus we cannot test whether there is a GBA trafficking defect with PGRN loss. Nevertheless, MDW41 labeling shows normal localization of active GBA in PGRN knockout fibroblasts (Fig. S4).

The relationship between PGRN and GBA is complicated by a series of interrelated factors. Two peptides derived from PSAP, saposin A and saposin C, are known activators of GBA (46, 53, 54). We have previously shown that PGRN and PSAP share a lysosomal co-trafficking relationship, wherein PGRN can carry PSAP to the lysosome via the receptor, sortilin, and PSAP can carry PGRN to the lysosome via the cation-independent mannose-6-phosphate receptor (CI-M6PR) or the low-density lipoprotein receptor-related protein 1 (LRP1) (30, 51, 52). Additionally, PGRN has recently been shown to bind and modulate the activity of the lysosomal protease, cathepsin D (CTSD), which is the major contributor to proteolytic PSAP processing. Because of these factors, it is possible that PGRN deficiency results in an alteration in PSAP processing and production of saposin peptides, thereby affecting GBA activation. However, PSAP, total saposin levels and saposin C levels do not appear to change in PGRN deficient tissue lysates (Fig. 3, Fig. S2), although we cannot rule out the possibility that saposin A levels might be affected.

We failed to detect any changes in GBA activity with the addition of recombinant PGRN or granulin peptides, but this does not entirely rule out the possibility of a direct activation of GBA by PGRN. This may require further optimization of the conditions for the assay.

Another possibility is that PGRN indirectly affects GBA activity by changing the lysosomal environment. PGRN deficiency has been reported to affect lysosomal pH (55), which could indirectly affect the activity of many lysosomal enzymes. Loss of PGRN also changes lipid contents of lysosomes (56), which could also indirectly affect GBA activity. While the exact mechanism of our findings is currently unknown, and more work needs to be performed to sift through the somewhat muddled relationships of the proteins involved, it is clear that PGRN deficiency leads to reduced GBA activity. This is significant, as lysosomal dysfunction is a commonality between NCL and FTLD with *GRN* mutation. It is possible that the decreased activity of multiple lysosomal hydrolases, including GBA and CTSD, accounts for lysosomal dysfunction with *GRN* mutations.

Mutations and polymorphism in the *GRN* gene have also been associated with Parkinson’s disease (PD)(57-60). In addition, reduced serum levels of PGRN were found to be associated with PD risk (61). Moreover, viral expression of PGRN was shown to protect midbrain dopaminergic neurons and improve the locomotor function in response to 1-methyl-4-phenyl-1,2,3,6-tetrahydropyridine (MPTP) treatment to mimic PD in mouse models (62). Interestingly, heterozygous mutations in *GBA* is one of the genetic determinants of PD(63) (64). Our work on the regulation of GBA activity by PGRN might provide a mechanistic explanation underlining the association of PGRN and PD.

## Conclusions

Our data support that PGRN deficiency leads to a reduction of GBA activity *in vivo.*

## Declarations

### Abbreviations

FTLD: frontotemporal lobar degeneration;
NCL: neuronal ceroid lipofuscinosis;
PGRN: progranulin;
GBA: glucocerebrosidase;
PSAP: prosaposin;
SAP: saposin;
SILAC: stable isotope labeling by amino acids in cell culture

## Ethical Approval and Consent to Participate

All applicable international, national, and/or institutional guidelines for the care and use of animals were followed. The work under animal protocol 2017-0056 is approved by the Institutional Animal Care and Use Committee at Cornell University.

## Consent for publication

All authors have given consent for publication.

## Availability of data and material

The datasets supporting the conclusions of this article are included within the article (and its additional files).

## Authors’ contributions

X.Z. and D.H.P collected all the data. M.D.P helped with activity assay and antibody testing. D.K and M.S. performed mass spec analysis. Y. S provided GBA deficient fibroblasts and tissue lysates. H. S. O provided MW probe. R. L.L. provided Cerezyme. F.H. performed the SILAC experiment, supervised the project and wrote the manuscript with D.H.P and X.Z. All authors read and approved the final manuscript.

## Acknowledgements

We would like to thank Dr. Haiyuan Yu for his gifts of cDNA; and Xiaochun Wu for technical assistance. This work is supported by funding to F. Hu from National Institute of Aging and National Institute of Neurological Disorders and Stroke (R01 NS095954) and President’s Council of Cornell Women Affinito-Stewart award. The authors declare no additional competing financial interests.

